# EcologicalNetworksDynamics.jl A Julia package to simulate the temporal dynamics of complex ecological networks

**DOI:** 10.1101/2024.03.20.585899

**Authors:** Ismaël Lajaaiti, Iago Bonnici, Sonia Kéfi, Hana Mayall, Alain Danet, Andrew P Beckerman, Thomas Malpas, Eva Delmas

## Abstract

1. Species interactions play a crucial role in shaping biodiversity, species coexistence, population dynamics, community stability and ecosystem functioning. Our understanding of the role of the diversity of species interactions driving these species, community and ecosystem features is limited because current approaches often focus only on trophic interactions. This is why a new modelling framework that includes a greater diversity of interactions between species is crucially needed.
2. We developed a modular, user-friendly, and extensible Julia package that delivers the core functionality of the bio-energetic food web model. Moreover, it embeds several ecological interaction types alongside the capacity to manipulate external drivers of ecological dynamics like temperature. These new features represent important processes known to influence biodiversity, coexistence, functioning and stability in natural communities. Specifically, they include: a) an explicit multiple nutrient intake model for producers, b) competition among producers, c) temperature dependence implemented via the Boltzmann-Arhennius rule, and d) the ability to model several non-trophic interactions including competition for space, plant facilitation, predator interference and refuge provisioning.
3. The inclusion of the various features provides users with the ability to ask questions about multiple simultaneous processes and stressor impacts, and thus develop theory relevant to real world scenarios facing complex ecological communities in the Anthropocene. It will allow researchers to quantify the relative importance of different mechanisms to stability and functioning of complex communities.
4. The package was build for theoreticians seeking to explore the effects of different types of species interactions on the dynamics of complex ecological communities, but also for empiricists seeking to confront their empirical findings with theoretical expectations. The package provides a straightforward framework to model explicitly complex ecological communities or provide tools to generate those communities from few parameters.

## Introduction

The bio-energetic model of food web dynamics has played a central role in the identification of the drivers of ecological community dynamics since its introduction by (Yodzis & Innes, 1992). Following its adaptation to complex trophic networks by (Brose *et al*., 2006b), it has been used to develop theory about a wide range of topics including stability of multispecies communities (Brose *et al*., 2006b), the dynamics of secondary extinctions (Brose *et al*., 2005) and the impacts of multiple threats on biodiversity (Binzer *et al*., 2012).

The reason for the success of the bio-energetic model is twofold. First, it facilitated the representation of complex mechanisms, while maintaining a low number of free parameters and simple mathematical formulation. This enabled to strategically address fundamental questions about several key patterns, theories and processes in ecology (Brose *et al*., 2006b; Martinez, 2020). Therefore, it became a fundamental bridge between several disconnected sets of theory and vital to defining theory about the impacts of global change on biodiversity and ecosystem function. Second, the bio-energetic model made it possible to investigate these ecological processes in truly complex communities. For the first time, theory was able to align with a diversity of empirical patterns in complex ecological systems, including evidence to explain the discrepancy between May’s predictions that large systems are unlikely to be stable (May, 1972; Brose *et al*., 2003) and empirical observations that large ecological communities are indeed observed in nature.

However, the bio-energetic model also suffers from limitations. It solely centres on trophic interactions, thereby overlooking the diverse array of other interactions observed in nature, such as competition and facilitation among species (Kéfi *et al*., 2012, 2015). Including these interactions into food webs has been demonstrated to have significant implications for community diversity, productivity, and stability (Miele *et al*., 2019; Kéfi *et al*., 2016).

Although, the bio-energetic model has been implemented in various programming languages and by numerous researchers (see for instance (Lurgi *et al*., 2014; Delmas *et al*., 2017; Sentis *et al*., 2021; Gauzens *et al*., 2023)), these frameworks do not immediately allow the possibility for users to consider multiple interaction types simultaneously. More generally, these frameworks often lack flexibility in the sense that it can be difficult to modify the model by specifying custom parameters or changing the modelling choices – for example, moving from logistic growth to explicit nutrient dynamics.

To meet this need, we developed a Julia package which aim to extend existing frameworks of the bio-energetic model by addressing aforementioned issues. Our implementation is fast, user-friendly, and flexible, allowing researchers to explore: explicit nutrient uptake by producers and thus exploitative competition (Brose, 2008), direct interference competition (Tilman, 1982), temperature dependence of ecological rates (Binzer *et al*., 2012, 2016), and non-trophic interactions (Kéfi *et al*., 2012; Miele *et al*., 2019). These additions come alongside the capacity to easily manipulate fundamental features of the bio-energetic framework such as species richness, network structure and complexity, body mass distributions, functional responses, as well as the scaling of metabolic, biomass production and foraging traits with body size.

Below, we begin by introducing the bio-energetic dynamical model. Subsequently, we delve into the new features added into our package. This is followed by an instructional guide on using the package, and finally, we provide a demonstration of each new feature through a series of use-case scenarios.

### The core bio-energetic dynamical model

We developed the Julia package, EcologicalNetworksDynamics, that implements two versions of the core, well established bio-energetic framework of Yodzis and Innes and others (Brose *et al*., 2006a; Binzer *et al*., 2016; Kéfi *et al*., 2012; Delmas *et al*., 2017), and integrates four additional frameworks that extend the model beyond simple trophic interactions. We first describe the core implementations of the bio-energetic model and then present the extensions that we developed.

We implemented two core approaches of the bio-energetic model frequently found in the literature: ‘bio-energetic’ and ‘classic’ They are distinguished by their scaling of time and the default consumption rate of a consumer according to the resource density (hereafter ‘functional response’). In the original ‘bio-energetic’ version, time is relative to the growth rate of a producer and the functional response is defined by typical Holling forms (see Eq. 1b). In the ‘classic’ version, time is absolute and the functional response is presented with time dependent attack rates and handling times.

### Bio-energetic version

Used in (Yodzis & Innes, 1992; Williams *et al*., 2007; Brose *et al*., 2006b) and implemented in (Delmas *et al*., 2017), time is expressed relative to the inverse of a producer intrinsic growth rate (*r*_*p*_). We write the model in Eq. 1.

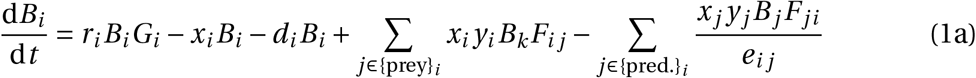

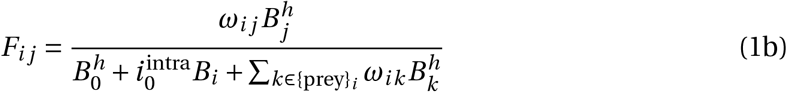

Where *B*_*i*_ is the biomass of the species *i, r*_*i*_ the intrinsic growth rate, *G*_*i*_ the normalized growth rate detailed further below, *x*_*i*_ the metabolic demand, *d*_*i*_ the natural mortality rate, {preys}_*i*_ and {pred.}_*i*_ respectively the ensembles of preys and predators of species *i, e*_*i j*_ the assimilation efficiency of species *i* feeding on prey *j*.

*F*_*i j*_ is the feeding rate of *i* feeding on *j* and depends on the half-saturation density *B*_0_ and *ω*_*i j*_ which weights the preference of predator *i* for prey *j*. By default, predators have the same preference for each of its prey, that is 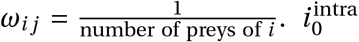 the intensity of intraspecific interference among consumers of a prey. *h* the hill exponent defining the shape (Type II or Type III) of the functional response: for (*h =* 1, 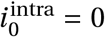) we recover the Holling type II functional response, and for (*h =* 2, 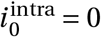) the type III.

### Classic version

Used in (Binzer *et al*., 2016) and also implemented in (Delmas *et al*., 2017), time has units that are absolute and the functional response depends on time dependent attack rates *a*_*i j*_ and handling times *h*_t,*i j*_. We write the model in Eq. 2.

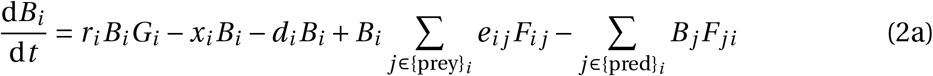

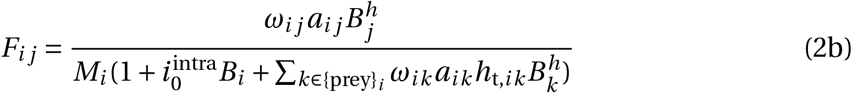

### Allometric scaling

A key feature of the bio-energetic model is the allometric scaling of metabolism, growth and foraging rates with body mass. This central role of allometry, reviewed in (Williams *et al*., 2007) makes for extremely efficient modelling of complex communities by reducing the number of free parameters and ‘automating’ the specifying of rates consistent with empirical data. Thus following (Yodzis & Innes, 1992) and all modern implementations of the bioenergetic model, we write the species’ intrinsic growth rate (*r*_*i*_), the metabolic demand (*x*_*i*_) and the natural mortality rate (*d*_*i*_) as function of their body-mass *M*_*i*_ (see Supporting Information Table S1). Following insights from (Brose *et al*., 2006a) and all other implementations of the model, we also assume that the consumer-resource body mass ratio (*Z*) can be any value but is constant across the network as the size structure of most empirical food webs is roughly constant. This allows the body mass of each species becomes a function of its trophic level (*T*_L_): 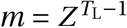, where we use the fractional trophic level after (Odum & Heald, 1975) using the calculation of (Pauly & Christensen, 1995).

### Extending the core model

We now present the new extensions of the bio-energetic model that we have implemented, namely: 1) producer competition, 2) nutrient dynamics, 3) temperature dependent biological rates, 4) non-trophic interactions, that all enables to model a diversity of complex ecological communities in various ecological context.

### Producer competition

By default, we model the producer growth (see Eq. 1a, 2a) as a logistic 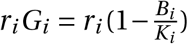, where *r*_*i*_ is the intrinsic growth rate of the producer *i* and *K*_*i*_ is the carrying capacity. However, the user can modify the relative strength of intra- versus interspecific competition. To do so, we modify the producer growth function to specify a competition coefficient in the numerator of the core logistic growth equation:

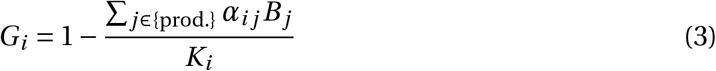

Two special cases emerge from this general form. First, if we assume no interspecific competition (for all *i* ≠ *j*, α_*i j*_ *=* 0) and that the intraspecific competition is set to unity (for all *i, α*_*ii*_ *=* 1), we recover to the logistic growth equation: 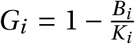. Second, in the case where intraspecific and interspecific competition are set to unity (that is, α_*i j*_ *=* 1.0 for all *i* and *j*), we assume that all producer species share a common pool of resources, so that 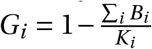 More generally, we can specify with this function multiple scenarios where the relative strength of intra-to interspecific competition can be evaluated.

### Nutrient uptake

In addition, the user can also change the default producer logistic growth for an explicit exploitative competition among producer species for nutrients. This modifies basal species’ growth rates *G*_*i*_ as follows (Brose *et al*., 2005; Brose, 2008):

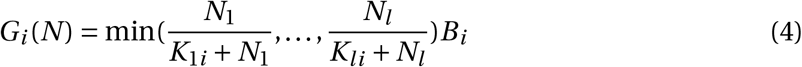

Where *N*_*l*_ is the concentration of nutrient *l* in the environment and *K*_*li*_ is the half saturation density of nutrient *l* for producer *i*. The nutrient-intake efficiency of producer *i* for nutrient *l* is then higher the lower its half-saturation density is.

The nutrient dynamics are determined by their respective supply (*S*_*l*_) and turnover rates (*D*_*l*_), their concentration in each producer (*c*_*li*_) as well as the producer’s half-saturation densities for the resource (*K*_*li*_) as shown in Eq. 5:

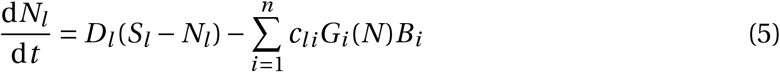

Default values for these parameters can be found in Supporting Information Table S3.

### Temperature dependence

Several researchers have integrated temperature dependence of the ecological rates in the bioenergetic model to explore questions, for example, about temperature interactions with productivity Binzer *et al*. (2016) and invasive species Sentis *et al*. (2021). Following these formulations, we implement in EcologicalNetworksDynamics the dependence of ecological rates with temperature using the exponential Boltzmann-Arrhenius relationship:

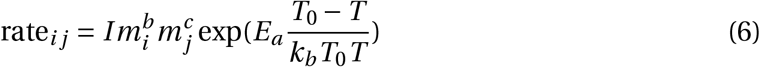

*I* is the allometric constant, *b* is the allometric exponent of species *i, c* is the allometric exponent of species *j, T* is the temperature of the system in Kelvin, *T*_0_ *=* 293.15 K is the normalisation temperature, *E*_*A*_ is the activation energy and *k*_*b*_ ≃ 8.617 *·* 10^*−*5^ VK^*−*1^ is the Boltzmann constant.

Species rates that can be scaled with temperature are the carrying capacities (*K*_*i*_), intrinsic growth rates (*r*_*i*_), metabolic rates (*x*_*i*_), attack rates (*a*_r,*i j*_) and handling times (*h*_t,*i j*_). As attack and handling time are specified, the temperature scaling of species rates rely on the ‘classic’ bio-energetic model (see previous section). Moreover, note that the attack rates and handling times depend both on resource species *i* and consumer species *j* (that is *c* ≠ 0 in Eq. 6), while the other rates only depend on the focal species *i* (*c =* 0). The default values of *I, b* and *c* for the different species rates can be found in Supporting Information Table S3.

### Non-trophic interactions

We implemented the possibility to model four non-trophic interactions that are ubiquitous and can have crucial effects on community dynamics and ecosystem functioning (Kéfi *et al*., 2016; Miele *et al*., 2019) : 1) competition for space between sessile species, 2) plant facilitation, 3) interspecific interference between predators and 4) provision of prey refuges from consumption. The effect of each non-trophic interaction can be translated in the model as a change in specific system parameters (Kéfi *et al*., 2012). Below, we detail how each non-trophic interaction is formally incorporated in the model. Non-trophic interactions are only implemented for the ‘classical’ bio-energetic model, consistently with (Kéfi *et al*., 2012; Miele *et al*., 2019).

### Competition for space

Competition for space can only occur between sessile species (mostly primary producers). We assume that two species competing for space will mutually decrease their net growth rate defined as the sum of the growth, consumption and metabolic loss terms of Eq. 2a:

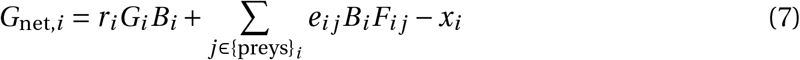

Then the effect of competition on the net growth rate is given by:

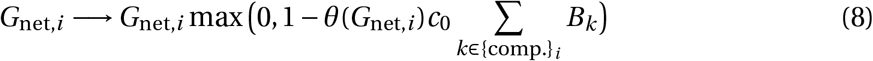

Where *c*_0_ is the intensity of the competition for space and {comp.}_*i*_ the ensemble of species competing with species *i*. *θ* is the Heaviside function (*θ*(*x*) *=* 1 if *x >* 0 and *θ*(*x*) *=* 0 otherwise). Therefore, the effect of competition on the net growth rate depends on its sign. If the net growth rate is positive, its value is reduced by the effect of competition and left unchanged otherwise. We ensure that a positive net growth cannot become negative by taking the maximum of zero and the multiplicative factor of the net growth rate.

### Plant facilitation

Producers experiencing recruitment facilitation have an increased intrinsic growth rate, following (Miele *et al*., 2019):

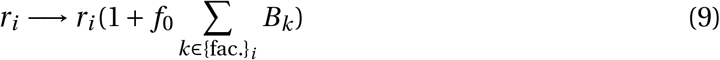

Where *f*_0_ is the intensity of the facilitation interaction, the larger *f*_0_ the more the producer growth rate is increased. Moreover, {fac.}_*i*_ is the ensemble of species, producers or not, facilitating the producer *i*.

### Interspecific interference between predators

Interference between predators can only occur between predators sharing at least one prey. We implement predator interference in the functional response by introducing a new term 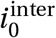 along with 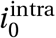

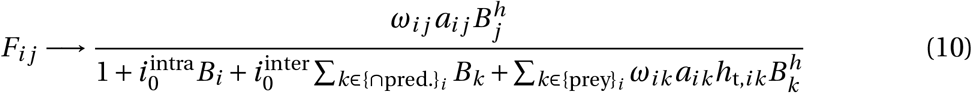

where 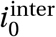 is the intensity of the interspecific interference and {*∩*pred.}_*i*_ the ensemble of predators sharing at least one prey with species *i*.

### Prey refuge

A refuge effect is translated as a reduction of the attack rates of the predators feeding on the prey *j* with a refuge. We model this by specifying the attack rate as a function that monotonically increases but plateaus with the strength of the refuge interactions. Refuge links occur from a sessile species (*i*) – which provides a refuge – toward a prey who benefits from the refuge (*j*).

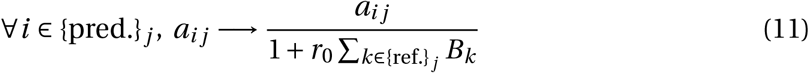

Where *r*_0_ is the intensity of the refuge effect and {ref.}_*j*_ the ensemble of species providing a refuge to species *j*.

### Basic usage

The EcologicalNetworksDynamics package enables users to simulate the dynamics of arbitrary complex ecological communities using an extended bio-energetic framework. Below, we describe the basic workflow of the package, *i*.*e*. how to: 1) create a network of interacting species, 2) define model parameters based on allometric relationships, and 3) simulate to produce a time series of biomass for all species. The package workflow is synthesized in Fig. 1. We then provide four use-cases for scenarios invoking competition, nutrient uptake, temperature dependence and non-trophic interactions. A guide to install Julia and the package can be found in the Supporting Information section S1.

**Figure 1.**
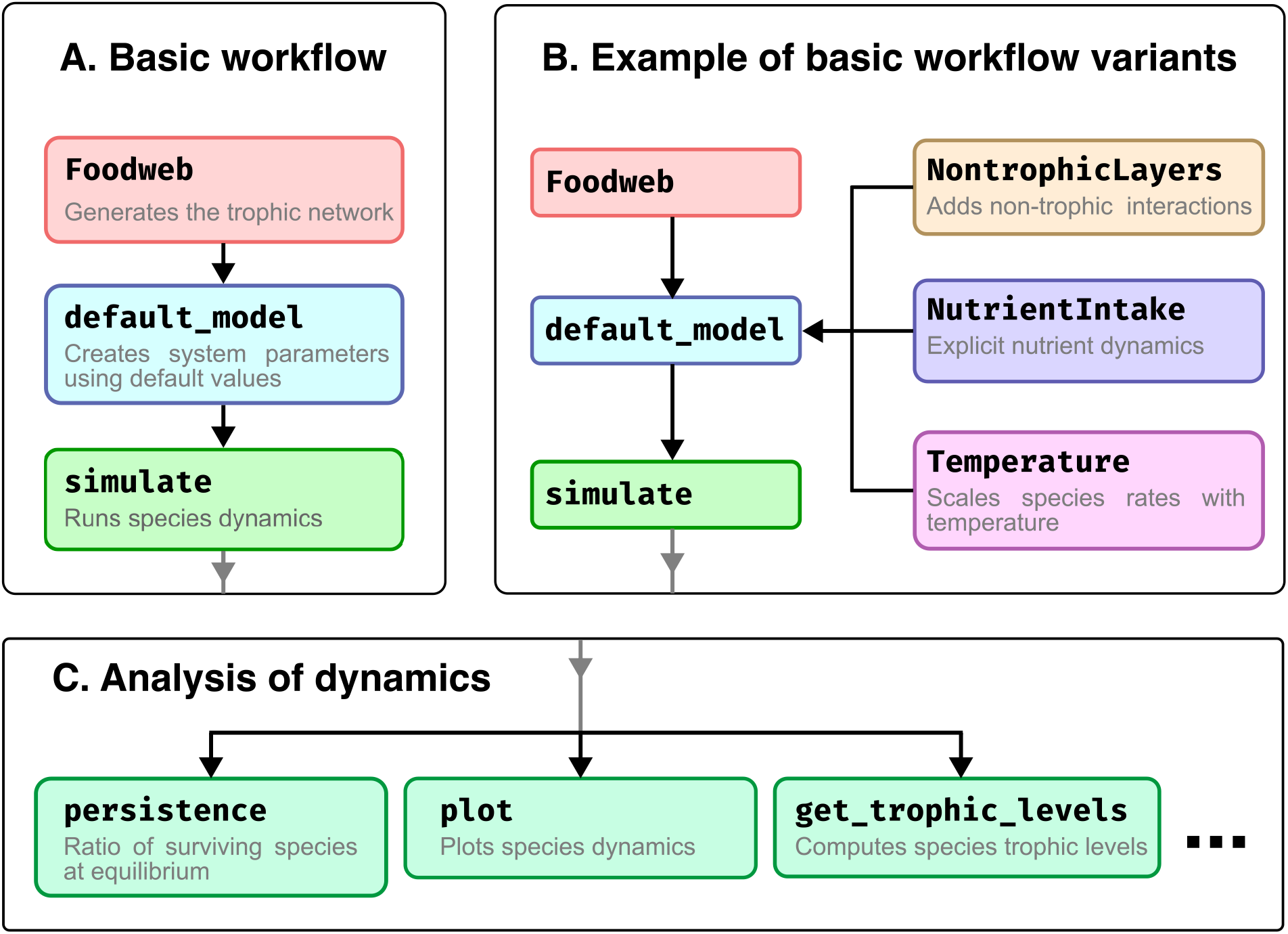
Overview of the package workflow: A) Most direct workflow predominantly using default package settings. B) Modified version of the basic workflow incorporating additional features to the model such as non-trophic interactions, nutrient dynamics, or temperature scaling of species biological rates. C) Examples of functions designed to analyse simulation outputs.

The following code simulates the dynamics of a primary producer (species 1) eaten by a consumer (species 2). The corresponding trophic network can be encoded into an adjacency matrix *A* (Eq. 12). EcologicalNetworksDynamics follows the convention of specifying that rows correspond to predator and columns to prey. This convention can differ from other implementations. In the adjacency matrix, 0s indicate the absence of trophic interactions while 1s indicate the presence of interactions. Moreover, EcologicalNetworksDynamics facilitates the use of several topological network generating models, including the cascade (Pimm, 1991) and niche models (Williams & Martinez, 2000).

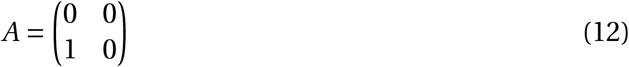

In this simple example, we specify that both species have individual body masses equal to 10, and initial population biomasses randomly drawn from the uniform distribution [0, 1]. By default, the ‘bio-energetic’ version of the model is used (Eq. 1) with default parameter values taken from the literature (see Supporting Information Table S**??** for default settings).

~~~
**using** EcologicalNetworksDynamics, Plots
foodweb = Foodweb([0 0; 1 0])
simple_model = default_model(foodweb, BodyMass([10, 10]))
B0 = rand(2) # Set initial biomasses for each species.
solution = simulate(simple_model, B0)
plot(solution)
~~~

We note that users can either provide a vector of body masses or set the predator-prey mass ratio (Z; PPMR -see Brose *et al*. (2006b)) which will distribute masses across trophic levels such that they scale exponentially with trophic levels. For example, to distribute body masses with a predator-prey mass ratio (PPMR) of 10 the model can be created as follows: `default_model(foodweb, BodyMass(Z = 10))`

Then, the `simulate` function calls under the hood the function `solve` from the Julia DifferentialEquations package (Rackauckas & Nie, 2017) which implements an adaptive time step solver. The choice of solver can be fully customized by the user and is detailed in the online documentation.

Lastly, the `plot` function of the Plots package (Christ *et al*., 2022) allow to directly visualize species trajectories. In addition, we provide several utility functions to analyse in more detail the result of the simulation. We showcase a few of these below and in Fig. 1:

~~~
total_biomass(solution) # Community biomass at each timestep.
total_biomass(solution[**end**]) # Final community biomass.
richness(solution) # Species richness at each timestep.
richness(solution[**end**]) # Final species richness.
~~~

### Use cases

Here, we showcase advanced and new features of the EcologicalNetworksDynamics package. The code to reproduce the examples can be found in the online documentation and in the associated GitHub repository (see Availability section). Figures from this section are made with the Julia plotting library `Makie` (Danisch & Krumbiegel, 2021).

### Paradox of enrichment

Classical theory associated with the paradox of enrichment (Rosenzweig, 1971) predicts a transition from stability to instability along a gradient of productivity defined by the carrying capacity *K* of the resource. This phenomenon appears in classic resource-consumer systems, where resources have density dependent growth and consumers feed with a Type II functional response. To reproduce this example, we define a trophic network with a single producer eaten by a consumer and run simulations for a range of producer carrying capacity *K*. For each simulation, we record the equilibrium biomass of each species. Results, recreating the paradox of enrichment by varying K, are shown in Fig. 2.

**Figure 2.**
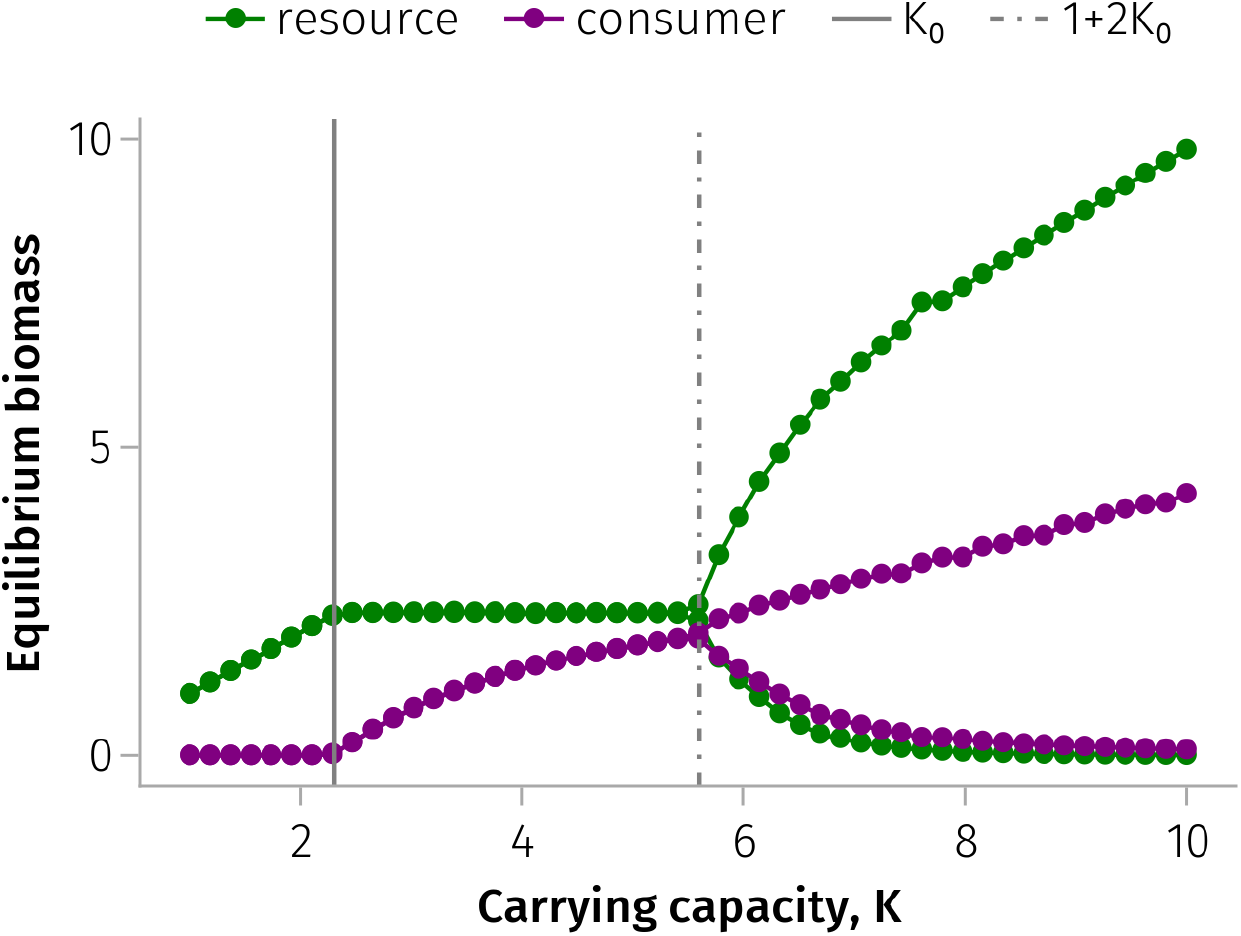
Orbit diagram of the consumer and resource biomasses (*B*) at equilibrium versus the resource carrying capacity (*K*). First, for *K ≤ K*_0_ there is not enough resource to sustain the consumer population, thus only the resource survives. Secondly, for *K*_0_ *≤ K ≤* 1 *+* 2*K*_0_ both species can coexist and the equilibrium attractor is a stable point. Moreover, we remark that in this region the consumer biomass increases with *K*, but not the resource one. Lastly, for *K ≥* 1 *+* 2*K*_0_ the system starts to oscillate and the amplitude of the limit cycle increases with *K* which can lead at some point to the species extinction. *K*_0_ is a critical carrying capacity, which mainly depends on the ratio of the consumer metabolic demand over its assimilation efficiency. This well-known pattern is referred to as the paradox of enrichment in the literature, and has been first described in 1971 by M. Rosenzweig (Rosenzweig, 1971).

### Producer competition

Coexistence theory predicts that stable coexistence is feasible when interspecific competition is lower than intraspecific competition (α_*i j*_ *<* α_*ii*_), while competitive exclusion arises when interspecific competition is higher than intraspecific one (α_*i j*_ *>* α_*ii*_). Competitive exclusion among primary producers then can trigger extinction cascades among consumers in food webs. However, it is known that increasing the consumer generalism, *i*.*e*. increasing the trophic connectance, can mitigate the destabilizing effect of a high interspecific competition. Here, we reproduce this result by measuring species persistence using a network of *S =* 20 species, three levels of connectance (complexity) and experimental manipulation of the interspecific producer competition coefficient α_*i j*_. To generate large realistic trophic networks, the package gives the possibility to create them from niche model (Williams & Martinez, 2000). For example, `Foodweb(:niche; S = 20, C = 0.1)` creates a network of 20 species with a trophic connectance. Results are show in Fig. 3.

**Figure 3.**
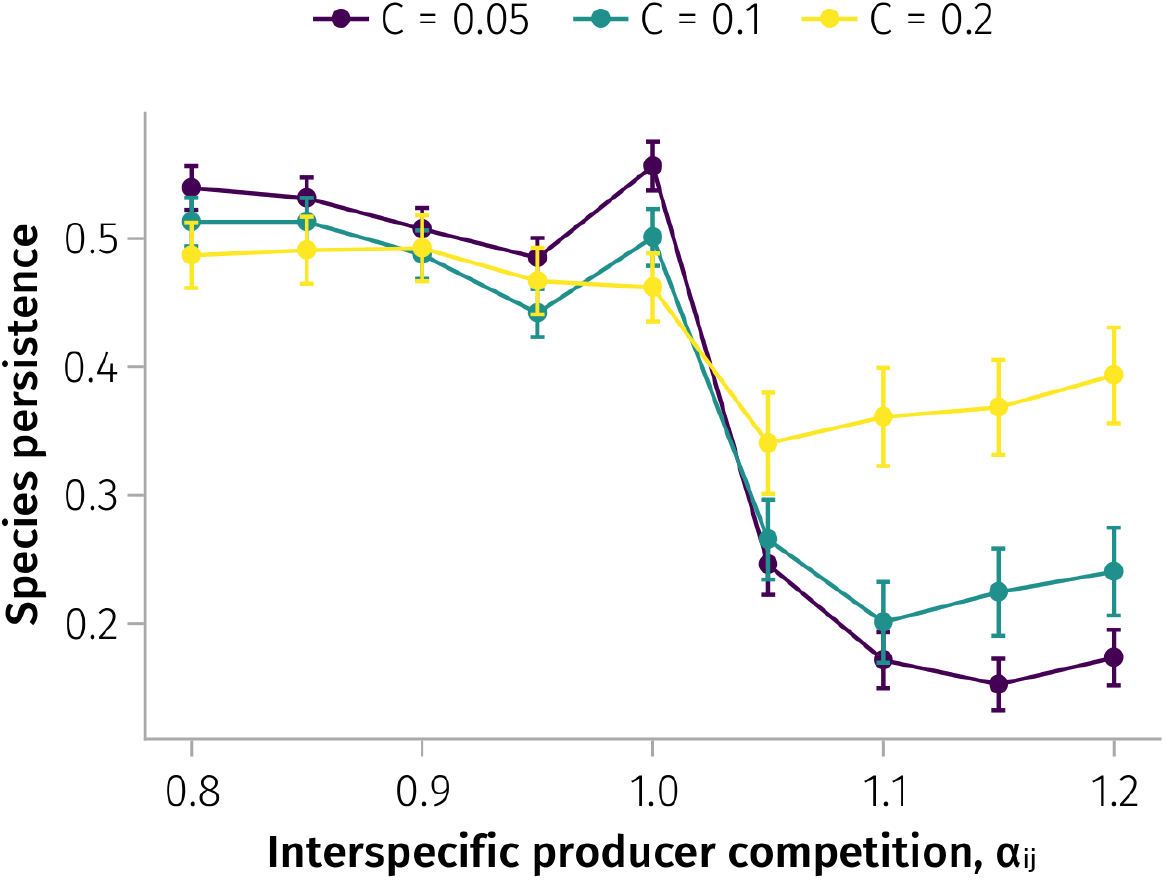
Species persistence along a gradient of interspecific competition strength among producers for three values of connectance. *S =* 20, *Z =* 100, *K =* 1.0, α_*ii*_ *=* 1. Species persistence drops when interspecific competition exceeds 1.0, i.e. when it becomes higher than intraspecific one. Higher values of connectance are associated with higher species persistence overall, especially when interspecific competition is higher than intraspecific one. Points display the average species persistence, the error bars display the 95% confidence intervals assuming a Normal distribution. This figure replicates results of (Delmas *et al*., 2017), Fig. 3.

### Nutrient uptake

Classical competition theory predicts that no two species can coexist on a single resource. However, when the number of resources is equal to or greater than the number of competitors, and there is a trade-off between growth rates and the ability to draw down each nutrient, coexistence can occur (Tilman, 1982; Huisman & Weissing, 1999).

To reproduce this classic theory, we define two networks: one with two producers competing for a single nutrient and one with two producer species competing for two nutrients. Results are shown in Fig. 4 where under the first scenario, competitive exclusion occurs and in the second, coexistence.

**Figure 4.**
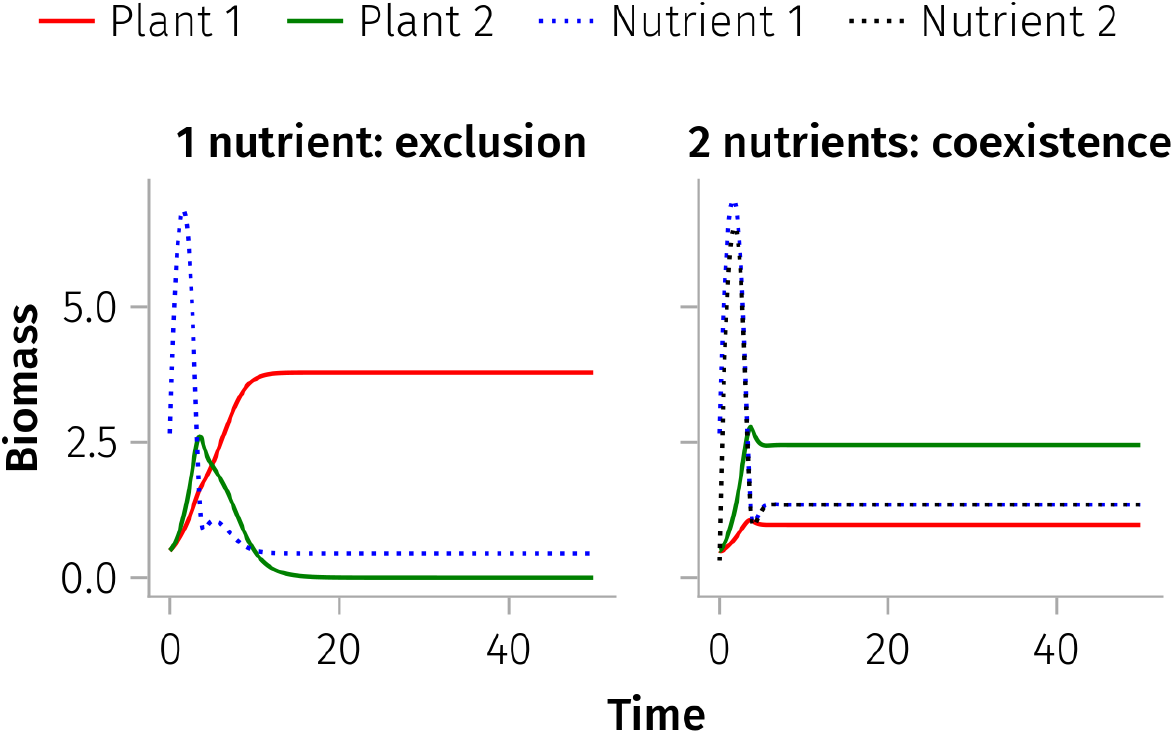
Biomass of two plant species sharing one (left) or two (right) nutrient resources through time. Following theory, the single shared resource leads to competitive exclusion because Plant 1 can draw down the resource to a lower equilibrium than plant species 2. In contrast, with two nutrients and a trade-off between growth rates and the ability to draw down each nutrient, coexistence can occur.

### Temperature dependence

Temperature dependence is by default turned off, but can be introduced with the component `Temperature`, that will redefine the temperature dependent rates using the exponential Boltzmann Arrhenius scaling.

Here we showcase the equilibrium dynamics of a simple 3-species trophic chain along a temperature gradient from 0 to 40°C (Fig. 5). Consumers feed with a Holling type-II (classic) functional response and carrying capacities (*K*_*i*_), intrinsic growth rates (*r*_*i*_), metabolic rates (*x*_*i*_), attack rates (*a*_r,*i j*_) and handling times (*h*_t,*i j*_) scale with temperature (see (Binzer *et al*., 2012) for a similar result with the bio-energetic functional response). We observe a transition from a basal trophic level only equilibrium at the producer carrying capacity at low temperatures and a state change around 17°C where coexistence arises, even though the producer remains at a low biomass.

**Figure 5.**
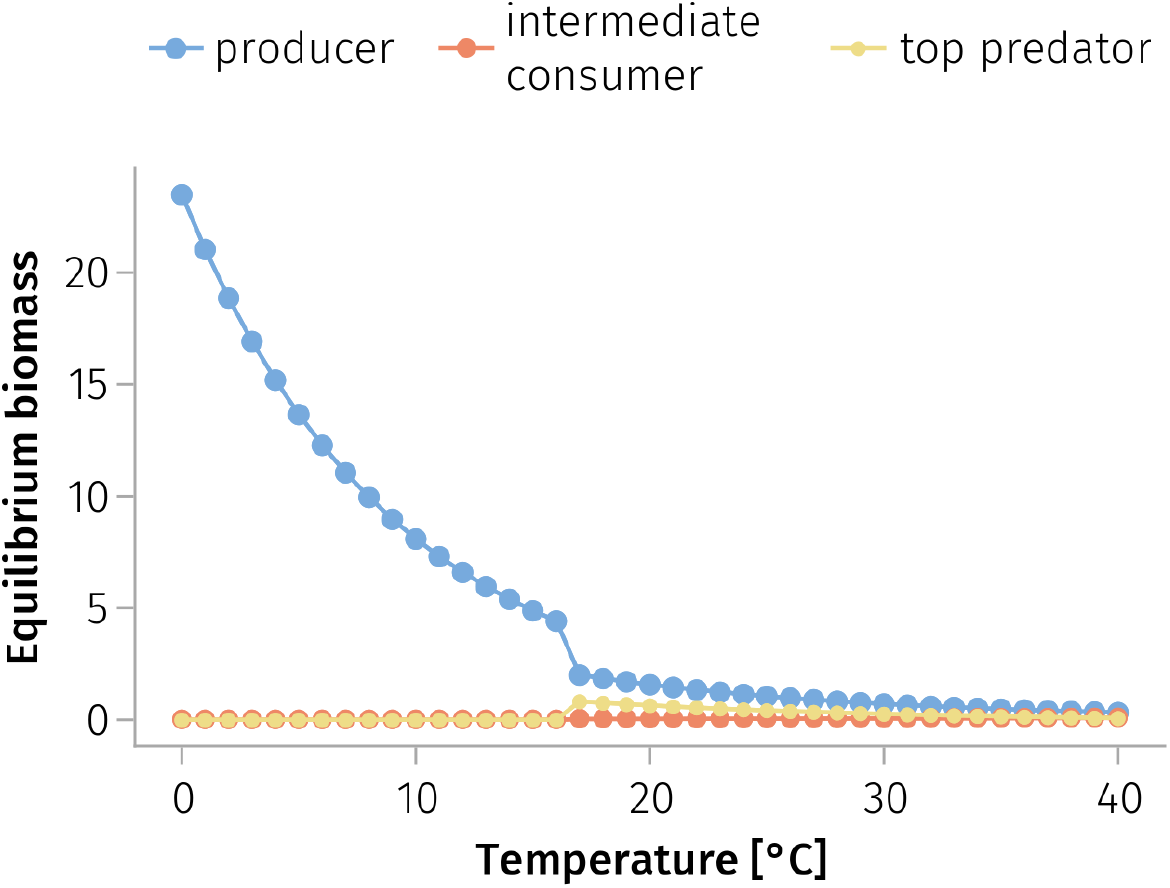
Temperature alters the equilibrium dynamic and coexistence of species in a tritrophic community. Each point records the equilibrium biomass of a species at a given temperature. At low temperatures, the basal producer species is the only surviving species, growing to the carrying capacity. Further along the temperature gradient, the system changes state with all three species coexisting at low biomass.

### Non-trophic interactions

Non-trophic interactions can be added to the food web by updating the food web with the `NonTrophicLayers` function. The detail and structure of each non-trophic interactions is specified through an adjacency matrix, a number of links or a connectance. In the two last cases, the non-trophic links are drawn randomly conditioned by few simple rules (*e*.*g*. plant facilitation links are always directed toward a plant).

We showcase here the effect of non-trophic interactions on species diversity. Specifically, we reproduce the results of (Miele *et al*., 2019) which demonstrates how the strength of four non-trophic interactions, considered separately, can impact on species diversity. (Miele *et al*., 2019) found that facilitation had a positive effect on species diversity, while refuge effect, interspecific predator interference and competition for space had a negative effect. We varied the intensity of each non-trophic interaction and recorded species diversity at equilibrium. We present the mean and 95% confidence interval for 50 replicate food webs of 50 species with connnetance = 0.06ś0.01. Results are shown in Fig. 6.

**Figure 6.**
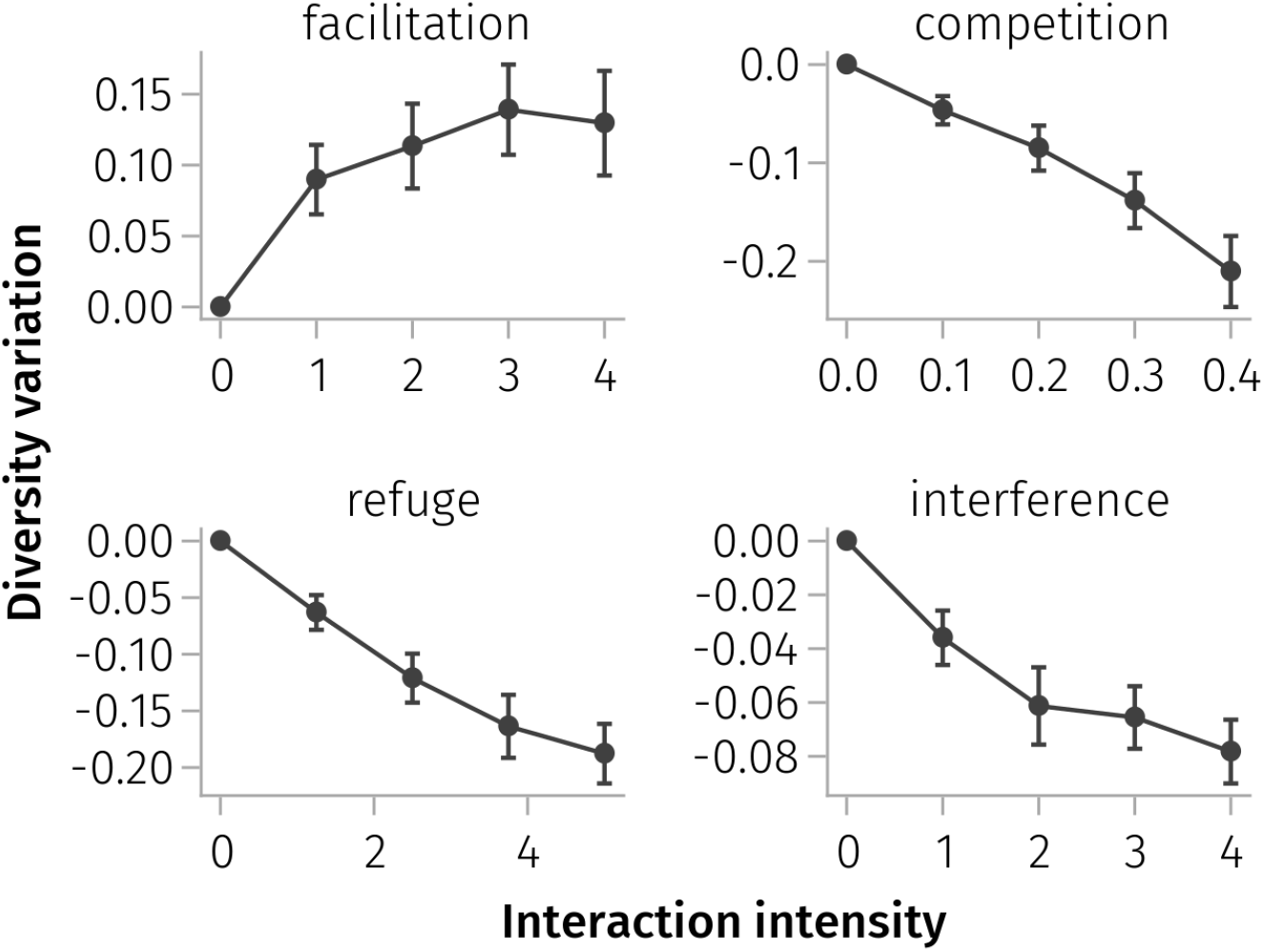
The relative variation in diversity as the intensity of the different non-trophic interactions is increased. Plant facilitation increases diversity while the three other non-trophic interactions decrease diversity. The points of zero intensity correspond to the reference, where there are only trophic interactions in the network. The relative variation in diversity is 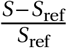, where *S* is the equilibrium richness of the multiplex network and *S*_ref_ of its reference trophic network, *i*.*e*. the multiplex network from which we have removed the non-trophic interactions. We start with community of *S*_init_ *=* 50 species and trophic links distributed with niche model for a connectance *C =* 0.06 *±* 0.01. Non-trophic interactions are drawn randomly given the rules of (Miele *et al*., 2019), with a connectance of *C*_NTI_ *=* 0.01. We dis-carded networks with loops or disconnected species. Error bars represent 95% confidence interval.

## Conclusion

We have presented `EcologicalNetworksDynamics`, a Julia package implementing the bioenergetic model with several extensions that include: 1) competition between producers; 2) an explicit nutrient uptake model for producers; 3) a temperature dependence of the model parameters; and 4) non-trophic interactions. The package is ideal for theoreticians seeking to explore the effects of different types of species interactions on the dynamics of complex ecological communities, but also for empiricists seeking to confront their empirical findings with theoretical expectations. It allows modelling communities from few parameters, while making possible for the user the possibility to customize the model by mixing interaction types and external drivers with ease. We believe that with this gain in flexibility over previous frameworks, our package will facilitate the exploration of and development of theory for a wide range of ecological scenarios. The outputs of the package can be exported in a language-neutral format (*e*.*g*. CSV), and thus be processed with other languages commonly used in Ecology for data analysis and visualisation, such as `R`. Critically, we believe this fast, extensible and open source code-base can help scientists and researchers avoid having to individually implement their own code, thus saving time and providing hopefully a common and extensible framework for the community.

### Package availability and development

The `EcologicalNetworksDynamics` package is available in the official Julia package registries and tested against the latest stable release of Julia (v1.10). Documentation, use cases above and source code are available at: https://github.com/BecksLab/EcologicalNetworksDynamics.jl/.

The package is released under GPL-3 licence and SemVer-compliant version number 0.2.0. The version number is meant to evolve, reflecting that the code authors did not unnecessarily constrain the user-facing library features (function names, syntax, workflow) to strict stability yet. We expect that future user feedback and additional features will be flexibly integrated into the package. If this evolution requires that the interface be modified in a non-retrocompatible way, then the changes will be rigorously documented in the package release notes, and user code will not break unless they explicitly upgrade the package as it eventually matures towards 1.0.0. In this respect, bug reports, improvement suggestions and contributions are very welcome under the form of ‘issues’ and ‘pull requests’ in the source repository.

## Acknowledgments

We thank Matthieu Fontaine for having extensively tested the package and reported many bugs to us. EcologicalNetworksDynamics benefited from the Montpellier Bioinformatics Biodiversity platform (MBB) supported by the LabEx CeMEB, an ANR “Investissements d’avenir” program (ANR-10-LABX-04-01). S.K. and I.L. were supported by the grant ANR-18-CE02-0010-01 of the French National Research Agency ANR (project EcoNet). A.P.B, A.D. and E.D. are supported by UKRI-NERC grants NE/T003502/1 and NE/S00-1395/1. H.M. and T.M. are supported by the UKRI-NERC Doctoral Training Programme ACCE with additional funding support from Unilever. We thank Timothée Poisot for critical feedback throughout the process and for initiating with Eva Delmas the original open source Julia package.

## Supporting Information

### A Getting started in Julia

The package is built in Julia and requires an up-to-date installation of this programming language. It is available for all operating systems (https://julialang.org/downloads/). Once Julia is installed, users can install our package by executing the following command in a Julia terminal (known as the REPL):

~~~
**using**
Pkg Pkg.add(“EcologicalNetworksDynamics”)
~~~

To make available the functionalities of the package, run:

~~~
**using** EcologicalNetworksDynamics
~~~

With this set-up, everyone should be able to execute and reproduce the examples presented in the following section. We note that VSCode, Atom and several other IDE environments offer methods for users to interact via scripting with Julia, instead of direct interaction through the REPL. We provide extensive documentation for use of the package on the associated GitHub repository and we advise the new users to go through the ‘Quick start’ page.

The documentation provides a user manual which explores the main features of the package one by one, and presents in depth several use cases including the examples given in this paper.Moreover, every public function of our package is documented.This help can be accessed directly from a Julia REPL with `?<function_name>` where `<function_name>` should be replaced by the name of the function of interest. For instance, you can type `?FoodWeb` to know how food webs can be generated.

### B Parameter tables and default settings

**Table 1.** Main parameter table. List of parameters within the package EcologicalNetworks-Dynamics, with their default values for producer and consumer species.

| Parameter | Symbol | Default producers | Default consumers | References |
| --- | --- | --- | --- | --- |
| Body mass | $M_i$ | 1 | 1 | |
| Intrinsic growth | $r_i$ | $M_i^{-0.25}$ | 0 | (Miele *et al.*, 2019) |
| Carrying capacity | $K_i$ | $M_i^{0.25}$ | ∅ | (Miele *et al.*, 2019) |
| Metabolism | $x_i$ | 0 | $0.314M_i^{-0.25}$ if invertebrates, $0.88M_i^{-0.25}$ vertebrates | (Delmas *et al.*, 2017)<br>(Miele *et al.*, 2019) |
| Maximum consumption | $y_i$ | 0 | 8 if invertebrates, 4 vertebrates | (Delmas *et al.*, 2017) |
| Mortality | $d_i$ | 0 | $0.0314M_i^{-0.25}$ | (Miele *et al.*, 2019) |
| Attack rate | $a_{ij}$ | ∅ | $50M_i^{0.45}M_j^{0.15}$ if $j$ is a Plant, $50M_i^{0.45}$ otherwise | (Miele *et al.*, 2019) |
| Handling time | $h_{t,ij}$ | ∅ | $0.3M_i^{-0.48}M_j^{-0.66}$ | (Miele *et al.*, 2019) |
| Preference | $\omega_{ij}$ | ∅ | $\frac{1}{\text{number of preys}}$ | (Delmas *et al.*, 2017)<br>(Miele *et al.*, 2019) |
| Efficiency | $e_{ij}$ | ∅ | 0.45 herbivores, 0.85 carnivores | (Delmas *et al.*, 2017)<br>(Miele *et al.*, 2019) |
| Half-saturation density | $B_0$ | ∅ | 0.5 | (Delmas *et al.*, 2017) |
| Hill exponent | $\omega_{ij}$ | ∅ | 2 | |
| Intraspecific competition | $i_0^{\text{intra}}$ | ∅ | 0 | |

**Table 2.** Temperature dependence of the different species rates. We consider a temperature dependence of the following rates: carrying capacity *K* (gm^*−*2^) (Meehan, 2006), intrinsic growth rate *r* (s^*−*1^) (Savage *et al*., 2004), metabolism (s^*−*1^) (Ehnes *et al*., 2011), attack rate (m^2^ s^*−*1^) and handling time (s) (Rall *et al*., 2012).

| | $K_i$ | $r_i$ | $x_i$ | $a_{r,ij}$ | $h_{t,ij}$ |
| --- | --- | --- | --- | --- | --- |
| $I$ | 3 | $\exp(-15.68)$ | $\exp(-16.54)$ | $\exp(-13.1)$ | $\exp(9.66)$ |
| $b$ | 0.28 | -0.25 | -0.31 | 0.25 | -0.45 |
| $c$ | 0 | 0 | 0 | -0.8 | 0.47 |
| $E_A$ | 0.71 | -0.84 | -0.69 | -0.38 | 0.26 |

**Table 3.** List of nutrient parameters and their default values. These values were taken from (Brose, 2008) Table 1.

| Nutrient parameter | Symbol | Default value |
| --- | --- | --- |
| Turnover | D | 0.25 |
| Concentration | C | 0.15 |
| Supply | S | 4 |
| Half-saturation | k | 0.1 |

## References

Binzer, A., Guill, C., Brose, U. & Rall, B.C. (2012) The dynamics of food chains under climate change and nutrient enrichment. Philosophical Transactions of the Royal Society B: Biological Sciences, 367, 2935–2944. Publisher: The Royal Society, 10.1098/RSTB.2012.0230.

Binzer, A., Guill, C., Rall, B.C. & Brose, U. (2016) Interactive effects of warming, eutrophication and size structure: Impacts on biodiversity and food-web structure. Global Change Biology, 22, 220–227. 10.1111/gcb.13086.

Brose, U. (2008) Complex food webs prevent competitive exclusion among producer species. Proceedings of the Royal Society B: Biological Sciences, 275, 2507–2514.

Brose, U., Berlow, E.L. & Martinez, N.D. (2005) Scaling up keystone effects from simple to complex ecological networks. Ecology Letters, 8, 1317–1325.

Brose, U., Jonsson, T., Berlow, E.L., Warren, P., Banasek-Richter, C., Bersier, L.F., Blanchard, J.L., Brey, T., Carpenter, S.R., Blandenier, M.F.C., Cushing, L., Dawah, H.A., Dell, T., Edwards, F., Harper-Smith, S., Jacob, U., Ledger, M.E., Martinez, N.D., Memmott, J., Mintenbeck, K., Pinnegar, J.K., Rall, B.C., Rayner, T.S., Reuman, D.C., Ruess, L., Ulrich, W., Williams, R.J., Woodward, G. & Cohen, J.E. (2006a) Consumerresource body-size relationships in natural food webs. Ecology, 87, 2411–2417. 10.1890/0012-9658(2006)87[2411:CBRINF]2.0.CO;2.

Brose, U., Williams, R.J. & Martinez, N.D. (2003) Comment on” foraging adaptation and the relationship between food-web complexity and stability”. Science, 301, 918–918.

Brose, U., Williams, R.J. & Martinez, N.D. (2006b) Allometric scaling enhances stability in complex food webs. Ecology Letters, 9, 1228–1236. 10.1111/j.1461-0248.2006.00978.x.

Christ, S., Schwabeneder, D., Rackauckas, C., Borregaard, M.K. & Breloff, T. (2022) Plots. jl–a user extendable plotting api for the julia programming language. arXiv preprint arXiv:220408775.

Danisch, S. & Krumbiegel, J. (2021) Makie.jl: Flexible high-performance data visualization for julia. Journal of Open Source Software, 6, 3349. 10.21105/joss.03349.

Delmas, E., Brose, U., Gravel, D., Stouffer, D.B. & Poisot, T. (2017) Simulations of biomass dynamics in community food webs. Methods in Ecology and Evolution, 8, 881–886. _print: 10.1111/2041-210X.12713, 10.1111/2041-210X.12713.

Ehnes, R.B., Rall, B.C. & Brose, U. (2011) Phylogenetic grouping, curvature and metabolic scaling in terrestrial invertebrates. Ecology Letters, 14, 993–1000. 10.1111/J.1461-0248.2011.01660.X.

Gauzens, B., Brose, U., Delmas, E. & Berti, E. (2023) Atnr: Allometric trophic network models in r. Methods in Ecology and Evolution, 14, 2766–2773.

Huisman, J. & Weissing, F.J. (1999) Biodiversity of plankton by species oscillations and chaos. Nature, 402, 407–410.

Kéfi, S., Berlow, E.L., Wieters, E.A., Joppa, L.N., Wood, S.A., Brose, U. & Navarrete, S.A. (2015) Network structure beyond food webs: mapping non-trophic and trophic interactions on chilean rocky shores. Ecology, 96, 291–303. 10.1890/13-1424.1.

Kéfi, S., Berlow, E.L., Wieters, E.A., Navarrete, S.A., Petchey, O.L., Wood, S.A., Boit, A., Joppa, L.N., Lafferty, K.D., Williams, R.J., Martinez, N.D., Menge, B.A., Blanchette, C.A., Iles, A.C. & Brose, U. (2012) More than a meal… integrating non-feeding interactions into food webs. Ecology Letters, 15, 291–300. 10.1111/j.1461-0248.2011.01732.x.

Kéfi, S., Miele, V., Wieters, E.A., Navarrete, S.A. & Berlow, E.L. (2016) How structured is the entangled bank? the surprisingly simple organization of multiplex ecological networks leads to increased persistence and resilience. PLoS biology, 14, e1002527.

Lurgi, M., Galiana, N., López, B.C., Joppa, L.N. & Montoya, J.M. (2014) Network complexity and species traits mediate the effects of biological invasions on dynamic food webs. Frontiers in Ecology and Evolution, 2, 36.

Martinez, N.D. (2020) Allometric trophic networks from individuals to socio-ecosystems: Consumer–resource theory of the ecological elephant in the room. Frontiers in Ecology and Evolution, 8, 92.

May, R.M. (1972) Will a large complex system be stable? Nature, 238, 413–414.

Meehan, T.D. (2006) Energy Use and Animal Abundance in Litter and Soil Communities. Ecology, 87, 1650–1658. 10.1890/0012-9658(2006)87[1650:EUAAAI]2.0.CO;2.

Miele, V., Guill, C., Ramos-Jiliberto, R. & Kéfi, S. (2019) Non-trophic interactions strengthen the diversityfunctioning relationship in an ecological bioenergetic network model. PLoS computational biology, 15, e1007269.

Odum, W.E. & Heald, E.J. (1975) The detritus-based food web of an estuarine mangrove community. Estuar Res Chem Biol Estuar Syst, 1, 265.

Pauly, D. & Christensen, V. (1995) Primary production required to sustain global fisheries. Nature, 374, 255–257.

Pimm, S.L. (1991) The balance of nature?: ecological issues in the conservation of species and communities. University of Chicago Press.

Rackauckas, C. & Nie, Q. (2017) Differentialequations.jl–a performant and feature-rich ecosystem for solving differential equations in julia. Journal of Open Research Software, 5.

Rall, B.C., Brose, U., Hartvig, M., Kalinkat, G., Schwarzmüller, F., Vucic-Pestic, O. & Petchey, O.L. (2012) Universal temperature and body-mass scaling of feeding rates. Philosophical Transactions of the Royal Society B: Biological Sciences, 367, 2923–2934. Publisher: The Royal Society, 10.1098/RSTB.2012.0242.

Rosenzweig, M.L. (1971) Paradox of enrichment: destabilization of exploitation ecosystems in ecological time. Science, 171, 385–387.

Savage, V.M., Gillooly, J.F., Brown, J.H., West, G.B. & Charnov, E.L. (2004) Effects of Body Size and Temperature on Population Growth. The American Naturalist, 163, 429–441. Publisher: The University of Chicago Press, 10.1086/381872.

Sentis, A., Montoya, J.M. & Lurgi, M. (2021) Warming indirectly increases invasion success in food webs. Proceedings of the Royal Society B, 288, 20202622.

Tilman, D. (1982) Resource Competition and Community Structure. Monographs in population biology. Princeton University Press.

Williams, R.J., Brose, U. & Martinez, N.D. (2007) Homage to Yodzis and Innes 1992: Scaling up feeding-based population dynamics to complex ecological networks. From Energetics to Ecosystems: The Dynamics and Structure of Ecological Systems, pp. 37–51. Springer.

Williams, R.J. & Martinez, N.D. (2000) Simple rules yield complex food webs. Nature, 404, 180–183. 10.1038/35004572.

Yodzis, P. & Innes, S. (1992) Body Size and Consumer-Resource Dynamics. The American Naturalist, 139, 1151–1175.

